# PBLR: an accurate single cell RNA-seq data imputation tool considering cell heterogeneity and prior expression level of dropouts

**DOI:** 10.1101/379883

**Authors:** Lihua Zhang, Shihua Zhang

## Abstract

Single-cell RNA sequencing (scRNA-seq) provides a powerful tool to determine precise expression patterns of tens of thousands of individual cells, decipher cell heterogeneity and cell subpopulations and so on. However, scRNA-seq data analysis remains challenging due to various technical noise, e.g., the presence of dropout events (i.e., excess zero counts). Taking account of cell heterogeneity and structural effect of expression on dropout rate, we propose a novel method named PBLR to accurately impute the dropouts of scRNA-seq data. PBLR is an effective tool to recover dropout events on both simulated and real scRNA-seq datasets, and can dramatically improve low-dimensional representation and recovery of gene-gene relationship masked by dropout events compared to several state-of-the-art methods. Moreover, PBLR also detect accurate and robust cell subpopulations automatically, shedding light its flexibility and generality for scRNA-seq data analysis.

## Introduction

RNA sequencing technology has provided us unprecedented opportunities to view the complex cellular systems such as disease or cancer^1^. However, conventional technology sequences millions of cells at a time and measures the profiling by average values, which leads to differences of cells being averaged. Recently, single cell RNA sequencing (scRNA-seq) has made a grand advance on throughput and resolution, which makes it a promising tool to study heterogeneous systems^2^. However, the quantity of mRNA in a single cell is so tiny that a million-fold amplification is often used. Therefore, only a fraction of transcripts may be captured during library preparation and a large amplification noise may be introduced during this stage. Thus, there is often a phenomenon named ‘dropout’ events in scRNA-seq data, in which a gene gets false zero or near zero values in some cells.

High ratios of ‘dropout’ may mislead further analyses such as low-dimensional representation, cell subpopulation identification and cellular developmental trajectory reconstruction. Many imputation methods designed for scRNA-seq have been developed in recent two years^3^. These imputation methods have various model assumptions, which model the missing value of a given gene in a specific cell according to the entries of its co-expressed genes and/or homogeneous cells. For example, MAGIC^4^ reconstructs the gene expression profile by a Markov affinity graph. scImpute^5^ firstly divides values into ‘dropout’ ones that need to be imputed and ‘confident’ ones that are not affected by dropout events with a mixture model. Then it imputes ‘dropout values’ with a non-negative least square model cell by cell. DrImpute^6^ adopts a ‘mean’ imputation strategy, which imputes zero values by averaging the corresponding ones in the same cluster. As the cluster number is often not known, it varies the number in certain range, and then obtains the final solution by averaging the values across this range. SAVER^7^, BISCUIT^8^, URSM^9^ are three Bayesian based methods. Among them, URSM is a supervised method needing cell labels in advance. BISCUIT and URSM usually take a relative long time to implement. Recent comprehensive comparison analyses^3^ indicate that scImpute and DrImpute may perform not so good on data with less collinearity, and SAVER and BISCUIT often imputes dropout with near zero values. Thus, an accurate and robust imputation method is still urgently needed.

Low-rank matrix recovery method approximating a low-rank matrix based on a few observable entries is a direct and powerful imputation strategy, which has shown promising performance in many fields^10–12^. It is essentially based on the correlation between rows and columns of the matrix. However, a recent study suggests that taking advantages of the presence of low-rank submatrices improves the performance than the traditional low-rank recovery^13^. As we know, scRNA-seq data exhibit large heterogeneity, indicting the existence of structured low-rank submatrices. Moreover, it has been demonstrated that gene expression levels have distinct effects on the dropout events^14^. Thus, integrating these characteristics into one framework to achieve effective expression recovery is of great potential.

To this end, we present a novel cell sub-Population based Bounded Low-Rank method (PBLR) for scRNA-seq data imputation, which well considers the cell heterogeneity and expression effects to dropouts. Applications to both simulated and real scRNA-seq data suggest that PBLR is an effective tool to recover transcriptomic level and dynamics masked by dropouts, improve low-dimensional representation, and restore the gene-gene co-expression relationship. Moreover, PBLR also detect accurate and robust cell subpopulations automatically, shedding light its flexibility and generality for scRNA-seq data analysis.

## Results

### Overview of PBLR

PBLR aims to impute zeros by taking in a raw scRNA-seq data *M* with *m* genes and *n* cells, where *M(i, j)* is the expression value of gene *i* in cell *j*. PBLR consists two components: (1) perform an ensemble clustering upon the scRNA-seq data of selected genes to determine *g* cell subpopulations as well as *g*+1 corresponding submatrices (*M^(k)^*, *k*=1, …, *g*+1) of the raw scRNA-seq data *M*, and (2) run a bounded low-rank matrix recovery method onto each submatrix *M^(k)^* (Fig. 1, see Methods for details). Specifically, PBLR first extracts a set of variably expressed genes and/or rare subpopulation specific genes as suggested by recent studies^15,16,17^. PBLR further employs non-negative matrix factorization (NMF) and Incomplete NMF to build a consensus matrix as the input of hierarchical clustering for determining final cell subpopulations and submatrices.

**Figure 1.**
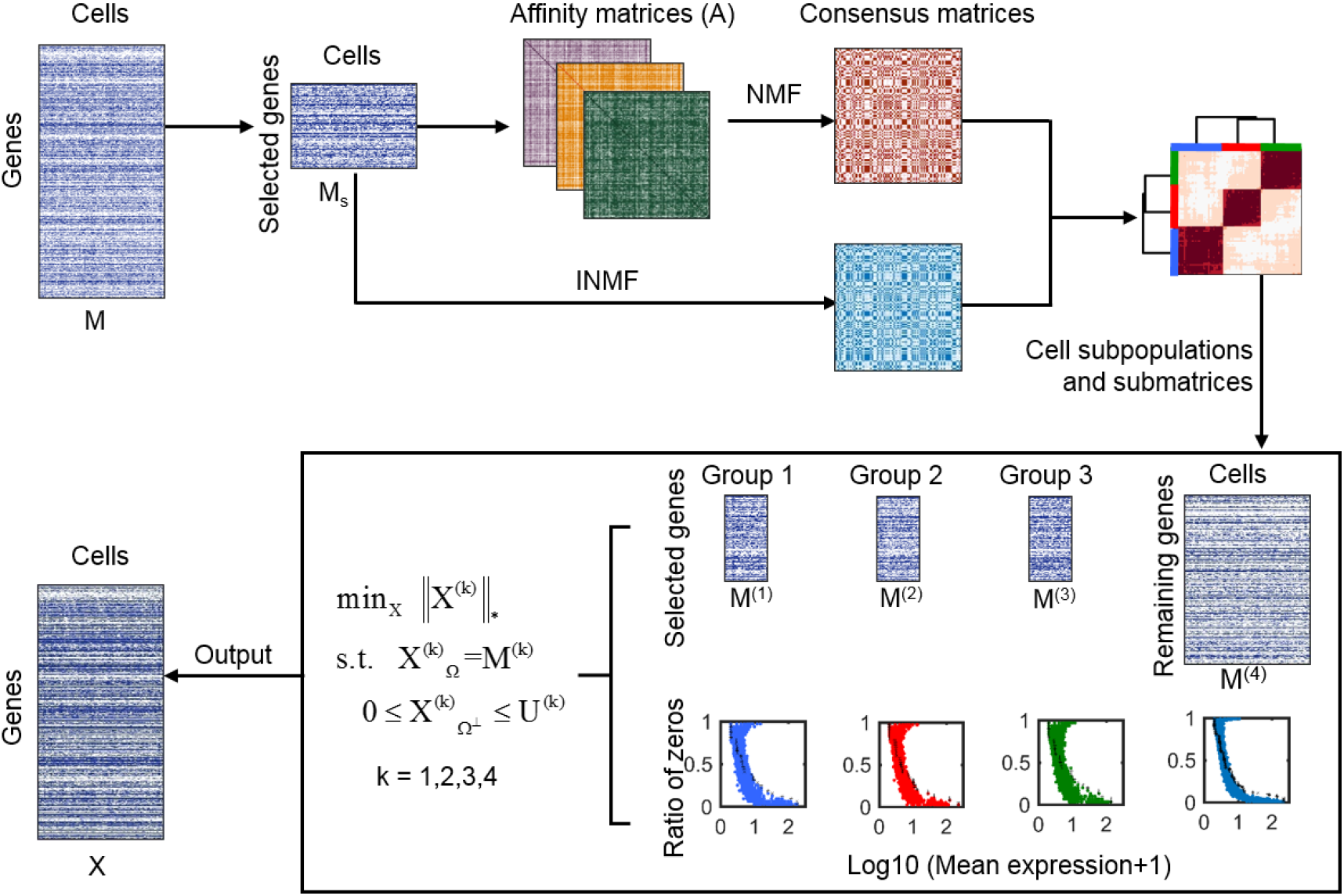
Overview of PBLR. Given a gene expression matrix *M* as input, PBLR outputs an imputed data matrix *X* with the same size as M. PBLR first extracts the data of selected high variable genes and computes three affinity matrices based on Pearson, Spearman and Cosine metrics respectively. Then, PBLR applies a symmetric non-negative matrix factorization (NMF) to the three affinity matrices of the sub-matrix of selected genes and incomplete NMF (INMF) to the sub-matrix to get the consensus matrices, respectively. PBLR further applies hierarchical clustering to the consensus matrix to infer cell subpopulations. Finally, PBLR estimates the expression upper boundary of the ‘dropout’ values, and recovers missed gene expressions by performing a bounded low-rank recovery model on each submatrix determined by cell subpopulations. In this diagram, there are three cell subpopulations. *M^(i)^, M^(2)^, M^(3)^* are the sub-matrices of each population of selected genes, *M^(4)^* is the sub-matrix of the remaining genes.

Let *X^(k)^* stand for the imputed data submatrix corresponding to the *k*-th submatrix *M^(k)^*. The low-rank recovery problem is formulated as follows,

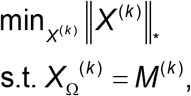

where Ω represents the so-called observed space in *M^(k)^* (i.e., the non-zero space), ||·||_*_ denotes the nuclear norm. Moreover, a recent study has shown that the probability of each gene’s dropout events varies across the expression magnitude, and there is a negative correlation relationship between the dropouts’ expression and the ratio of zeros^14^. Thus, the upper boundary of dropout values for a gene could be estimated in advance based on its observed expression level in other cells, which will improve the recovery accuracy (Supplementary Fig. 1). Therefore, PBLR introduces upper boundaries for unobserved variables, and then the bounded low-rank matrix recovery model is formulated into the following optimization problem,

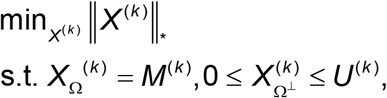

where Ω^⊥^ represents the unobserved space or say zero space, *U^(k)^* is a matrix in which each row denotes the upper boundary of a gene expression in the *k*-th submatrix *M^(k)^* (see Methods for details). This model is optimized by an efficient alternating Direction Method of Multipliers (ADMM) algorithm^18,19^. PBLR obtains the final imputed matrix *X* by merging these imputed submatrices *X^(k)^*.

### PBLR improves imputation accuracy of the low-rank discovery method by considering the cell heterogeneity and prior expression level of dropouts

Compared to a typical low-rank discovery model (LR), PBLR considers the structured characteristics of raw data and expression distribution reflected by the observed data to account for both cell- and gene-specific features of scRNA-seq data. To demonstrate the superior performance of these two key components, we used Splatter^20^ to generate synthetic dataset 1 with dropouts including three sub-populations (Supplementary Table 1). Visualization by principal component analysis (PCA) on the full data (data without dropouts) clearly shows three separated subpopulations or clusters. However, the clusters are confounded on the raw data due to the existence of dropouts (Fig. 2a). We applied LR to impute the raw data, and revealed mixed clusters (subpopulations) in the PCA space. Interestingly, performing LR on the inferred sub-matrices determined by cell sub-populations (denoted as PLR) can well separate them with more disperse clusters than those in the full data. However, it tends to over-estimate the expression of low-expressed genes compared to the real expression levels (Fig. 2b). Based on PLR, by further taking expression upper boundary into account, PBLR imputed data shows well separated clusters and more consistent distributions to the full data in the low-dimensional space (Fig. 2a) as well as more reasonable expression-to-dropout relationships (Fig. 2b). As expected, compared to LR and PLR, PBLR gives more accurate imputed values (Fig. 2c and d) in terms of sum of squared error (SSE) and Pearson correlation coefficients (PCC) (Methods).

**Figure 2.**
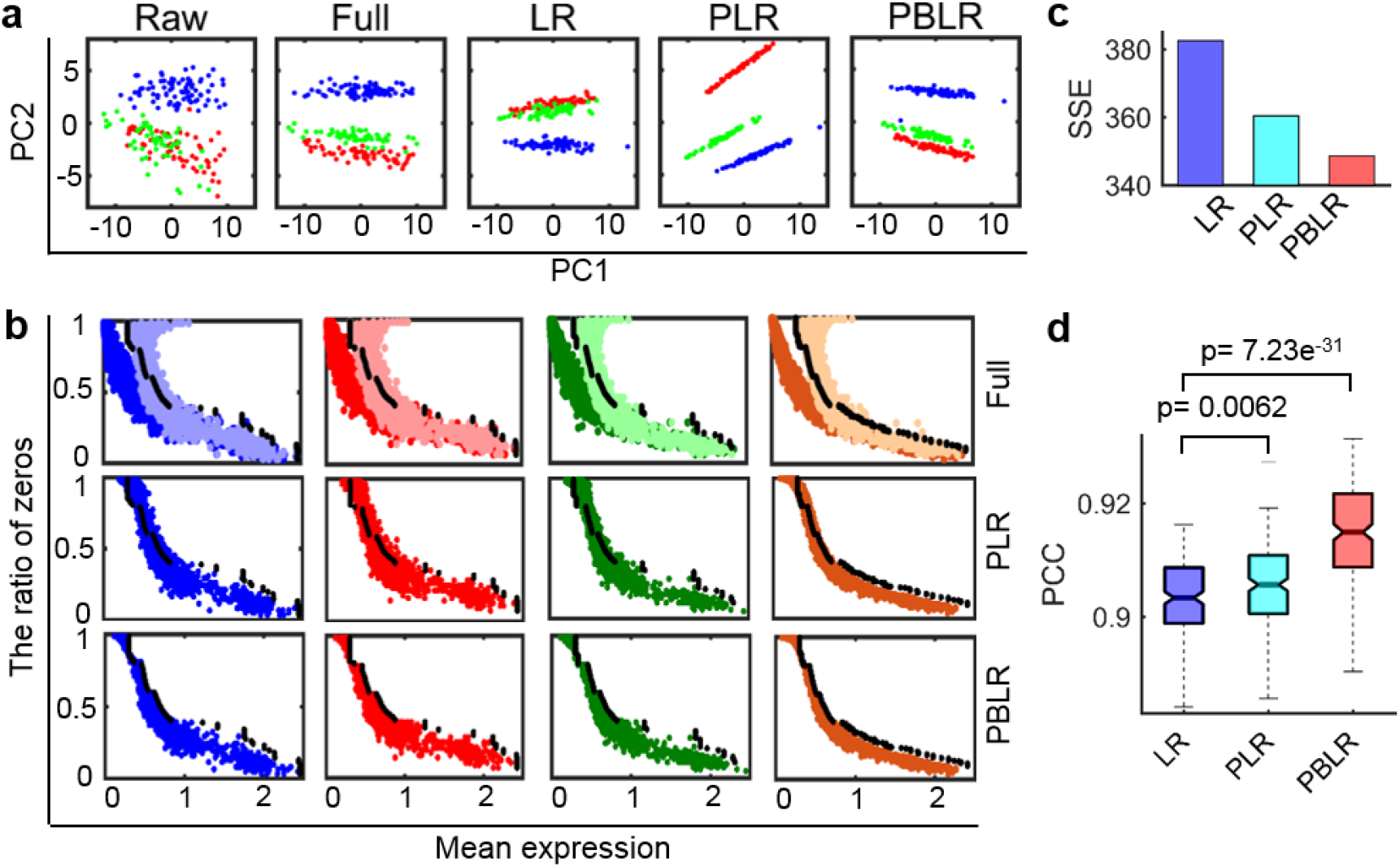
Comparison of PBLR with LR and BLR. **(a)** PCA visualization of the raw data, full data as well as imputed ones by LR, PLR and PBLR, respectively. LR represents the typical low-rank matrix recovery method, PLR indicates the population-based LR method. **(b)** Scatter plots of each gene with x axis representing log-transformed mean gene expression value and y axis representing the ratio of zeros across cells of each group. The top row shows distribution of real values of full data in the zero space (dark color) and non-zero space (light color) respectively for each sub-matrix. The middle and bottom rows show that of imputed values by PLR and PBLR for each sub-matrix respectively. Dots in different colors stand for imputed values of each sub-matrix in the zero space. The black dots represent the upper boundary. **(c)** SSE computed between the full data and the imputed ones by LR, PLR and PBLR respectively. **(d)** PCC computed for all single cell pair between the full data and the imputed ones by LR, PLR and PBLR respectively. *P*-value is computed by one-side Wilcoxon rank-sum test.

### PBLR recovers dropouts with superior accuracy compared to two competing methods

To demonstrate the effectiveness of PBLR, we compared it with two competing imputation methods (i.e., scImpute^5^ and SAVER^7^) in two aspects: the gene expression recovery and the effects on low-dimensional representation. To show performance of the imputation methods with respect to different dropout rates, we simulated synthetic dataset 2 with the shape parameter of dropout logistic function (ds) equaling −0.20, −0.15, −0.1, −0.05 corresponding to different ratios of zeros varying from 0.6 to 0.71 (Supplementary Table 1). We divided the entries of raw expression data into zero space and non-zero space. In the zero space, the imputed values of SAVER are much smaller than the real ones. While scImpute gives much larger fluctuations than PBLR (with ds=-0.05 as an example in Fig. 3a). Thus, PBLR recovers more similar values to the real ones than scImpute and SAVER. In the non-zero space, scImpute treat many moderate expression values as dropouts and imputes them by larger or smaller values than the real ones (Fig. 3a). Moreover, we also evaluated the imputation performance of scImpute, SAVER and PBLR in terms of SEE and PCC (Fig. 3b and c). As expected, the SSE values increase and PCC values decrease with the increase of the rates of zeros for these imputation methods. All these imputed data improve the performance of SSE and PCC relative to the raw data. Attractively, PBLR shows the smallest SSE values and largest PCC values compared to scImpute and SAVER. Visualization by the first two t-SNE components show that three real cell subpopulations in full data are mixed together as the existence of large amounts of zeros in raw data. SAVER plays almost no effect on the raw data. scImpute leads to three fictitious cell subpopulations in the t-SNE space, and shows improved performance in a dataset with a relative larger number of genes (Supplementary Fig. 2). However, the cell clusters can be well separated after applying PBLR. In summary, PBLR shows a strong ability in recovering dropouts compared to scImpute and SAVER on synthetic dataset 2 with various dropout rates (Fig. 3) and synthetic dataset 3 with a relative larger scale (Supplementary Note and Supplementary Fig. 3).

**Figure 3.**
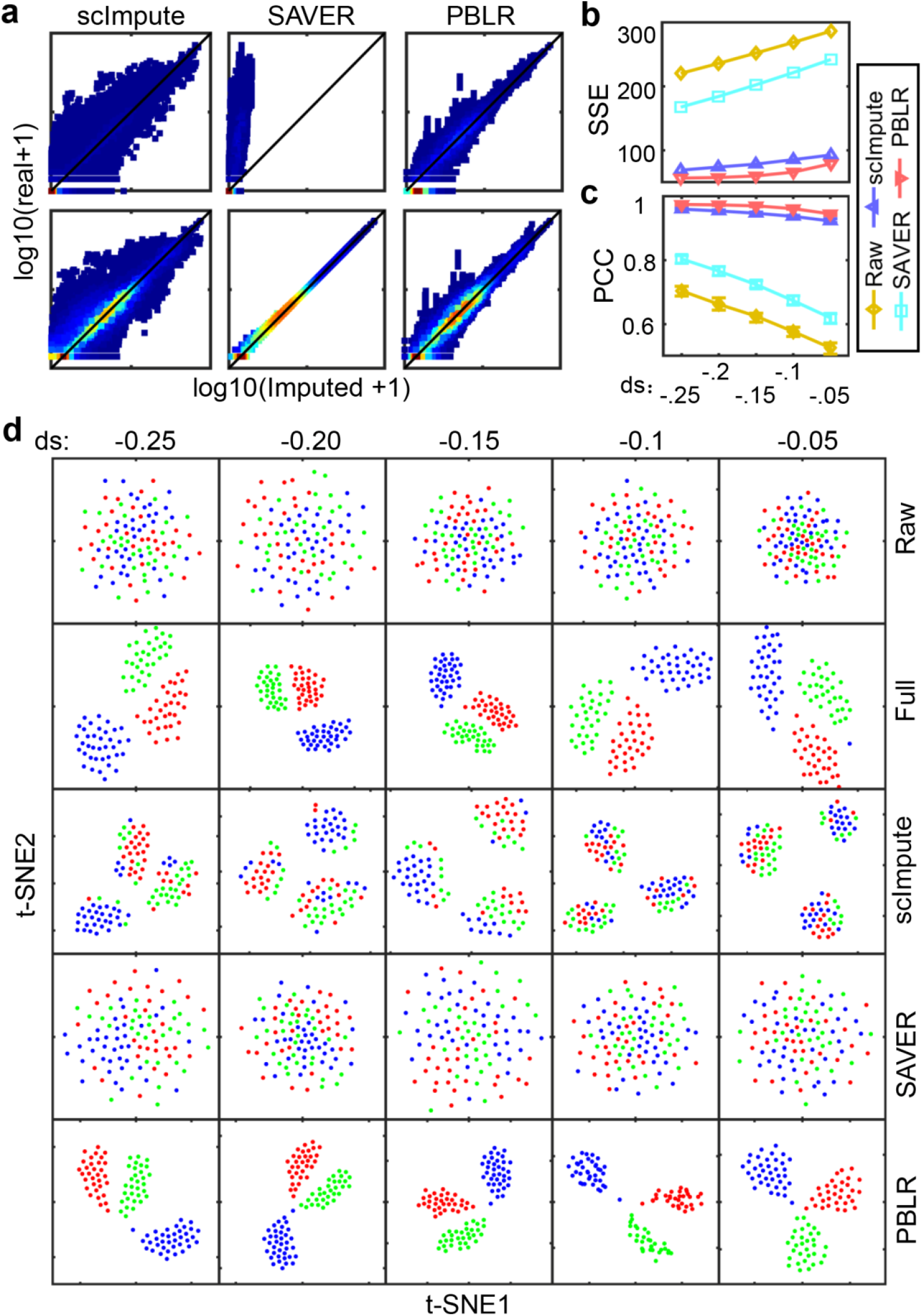
Imputation performance of scImpute, SAVER and PBLR on synthetic dataset 2 with various dropout rates. **(a)** Density plot of the imputed values versus true ones in the zero space (top) and the non-zero space (bottom), respectively. **(b)** Sum of squared error (SSE) values computed between the full data and the raw data as well as imputed ones respectively. **(c)** PCC values computed between the full data and the raw data as well as imputed one by scImpute, SAVER and PBLR respectively. **(d)** Visualization of cells by the first two t-SNE components on the raw data and imputed ones by scImpute, SAVER and PBLR respectively. Each column represents data with one dropout rate. **ds** means the parameter of dropout.shape in splatter package, which controls the ratio of zeros and larger value represents larger ratio of zeros in the data.

### PBLR captures precise expression dynamics during human early embryo development

We used scRNA-seq data consisting of 88 cells from seven stages (from oocytes to blastocyst) in human early embryos (HEE)^21^ to show whether the imputation values have biological meaning or not. First, PBLR accurately reveals the similarity of cells in each stage and cells in consecutive stages, and clearly capture the cell subpopulations (Fig. 4a). More interestingly, it identifies two cell subpopulations (denoted by G1 and G2) at the late blastocyst stage, in which various marker genes are considered to be expressed. It has been reported that *CDX2* is highly expressed in trophectoderm (TE), *SOX2, NANOG* and *KLF4* are highly expressed in epiblast (EPI) but lowly expressed in primitive endoderm (PE), and *FGFR4* and *CLDN3* are highly expressed in primitive endoderm (PE)^21^. Based on the marker genes mentioned above, we can see that TE cells and PE cells are enriched in G1 group, while EPI cells are enriched in G2 group (Fig. 4b). Some zero values of these marker genes are imputed by scImpute, SAVER and PBLR. For example, *CDX2* is imputed by scImpute and SAVER. And *SOX2* is imputed by PBLR (Fig. 4b). At blastocyte stage, two critical segregations take place: the segregations of cells into inner cell mass (ICM) and TE cells, and further differentiation of ICM cells into EPI and PE. Therefore, the expression of *CDX2* and *SOX2* exhibits negative correlation relationship, while that of *NANOG* and *SOX2* shows positive correlation relationship. After imputation, scImpute, SAVER and PBLR enhance the relationship of these two pairs of marker genes in different degree (Fig. 4c, Supplementary Fig. 4a). Attractively, PBLR significantly decreases the correlation between *CDX2* and *SOX2* from −0.37 to −0.53, and increases the correlation between *NANOG* and *SOX2* from 0.44 to 0.65. In addition to test these marker genes, we downloaded TE, EPI, and PE enriched marker genes (Supplementary Table 2) from a previous study^21^. Our results demonstrate that scImpute and SAVER slightly enhance the gene-gene correlation relationships (p-value > 0.05, one-sided Wilcoxon rank-sum test), however, PBLR is able to significantly enhance them including both positive and negative correlations (Fig. 4d), indicating its effectiveness in capturing the subtle expression relationship.

**Figure 4.**
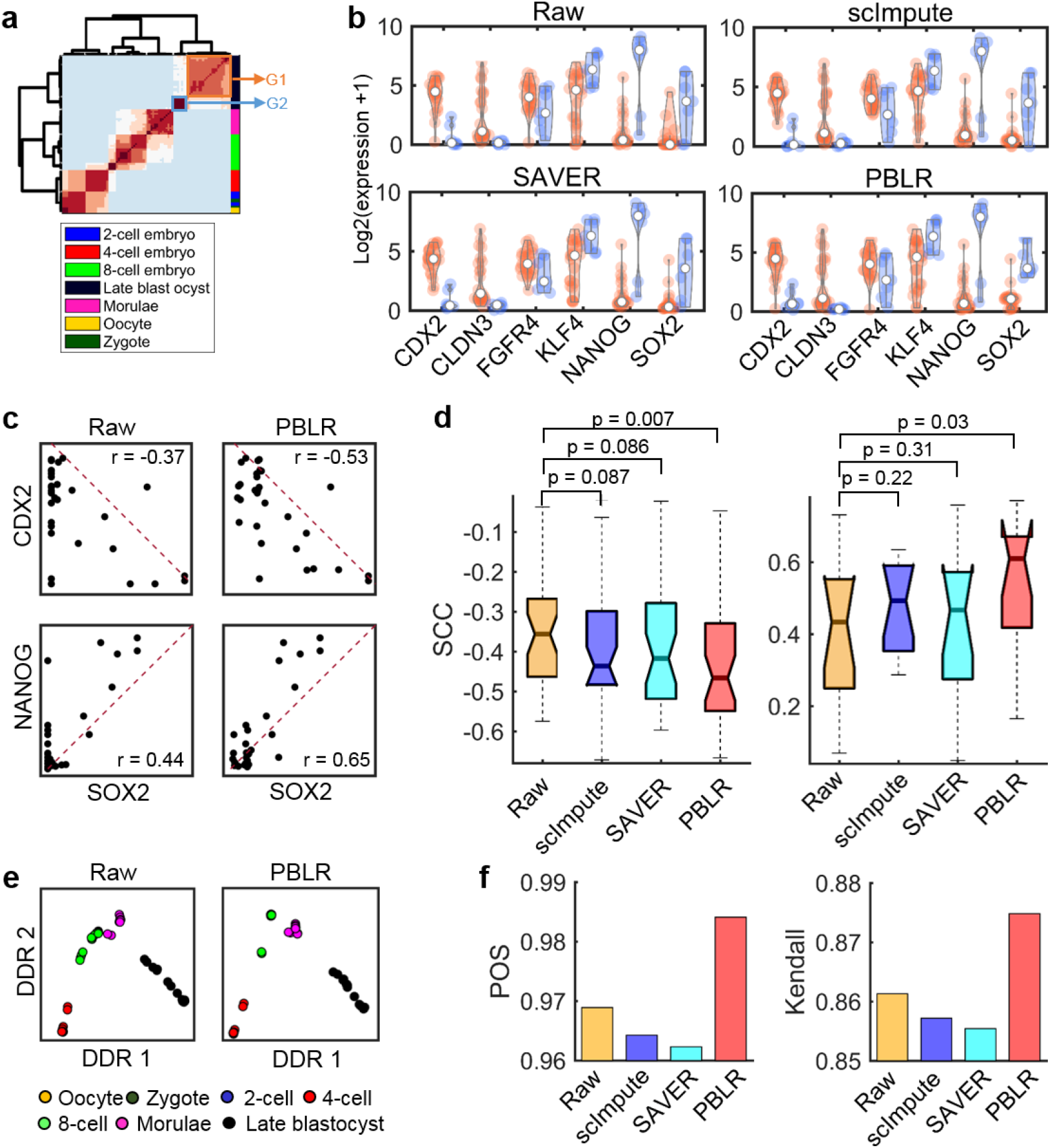
Marker gene expression patterns revealed on the real data from human early embryos development. **(a)** Hierarchical clustering on the consensus matrix obtained by PBLR. Experimental stage of each cell is indicated by different colors on the right. The late blastocyst cells are divided into two groups G1 and G2. **(b)** Violin-plot of gene expression values of marker genes in G1 (orange), G2 (light blue) groups. **(c)** Scatter plots of marker genes’ expression in the raw and imputed data by PBLR respectively. The corresponding SCC of expression values in the late blastocyst cells is shown on the top. **(d)** Comparison of SCC values of gene pairs from any two enriched gene sets for TE, EPI and PE (top), and gene pairs within EPI specific gene set (bottom) between imputed data and raw data. x-axis indicates the SCC values of the raw data and y-axis indicates the SCC values after applying scImpute, SAVER and PBLR respectively. Each dot represents a gene pair. *P*-values are computed by one-sided Wilcoxon rank-sum test. **(e)** Scatter plots of the first two discriminative dimensions inferred by Monocle 2. Each dot represents one cell. **(f)** Bar plots of POS scores and Kendall’s rank correlation coefficients after applying Monocle 2 to the raw and imputed data by scImpute, SAVER and PBLR, respectively.

Finally, we applied Monocle 2^22^ to the human early embryo development (HEE dataset) and the reprogramming from mouse embryonic fibroblasts (MEFs) to induce neuronal (iN) cells (MEF dataset)^23^, and imputed them to test whether PBLR can recover gene expression temporal dynamics (Fig. 4e, Supplementary Figs. 4b and 5). The major developmental trajectory can be detected on both raw data and PBLR imputed data visually (Fig. 4e, Supplementary Fig. 4b). We can clearly see that Morulae stage cells are in more compact cluster in the first two discriminative dimensions inferred by Monocle 2 (Fig. 4e). PBLR improves the inference performance distinctly compared to that of raw data, scImpute and SAVER imputed ones by applying Monocle 2 in terms of pseudotime order score (POS) and Kendall’s rank correlation (Fig. 4f, Supplementary Fig. 5b).

### PBLR improves the identification of cell subpopulations on real scRNA-seq datasets

As we can see that PBLR can not only impute missing values, but also reveal cell subpopulations directly from the raw data. Several powerful clustering methods specially designed for scRNA-seq have been proposed^24–26^. We applied PBLR to five real scRNA-seq datasets and compared it with SC3^24^, Seurat^25^, SIMLR^26^ and k-means on the first two t-SNE dimensions. The ratios of zeros of these datasets vary from 60.5% to 90.2% (Supplementary Table 3). Generally, among these clustering methods, PBLR and SC3 performs better and stable than other methods. PBLR exhibits the highest accuracy than other clustering methods on raw data except for Darmanis dataset (Fig. 5a), indicating its distinct superiority to competing methods.

**Figure 5.**
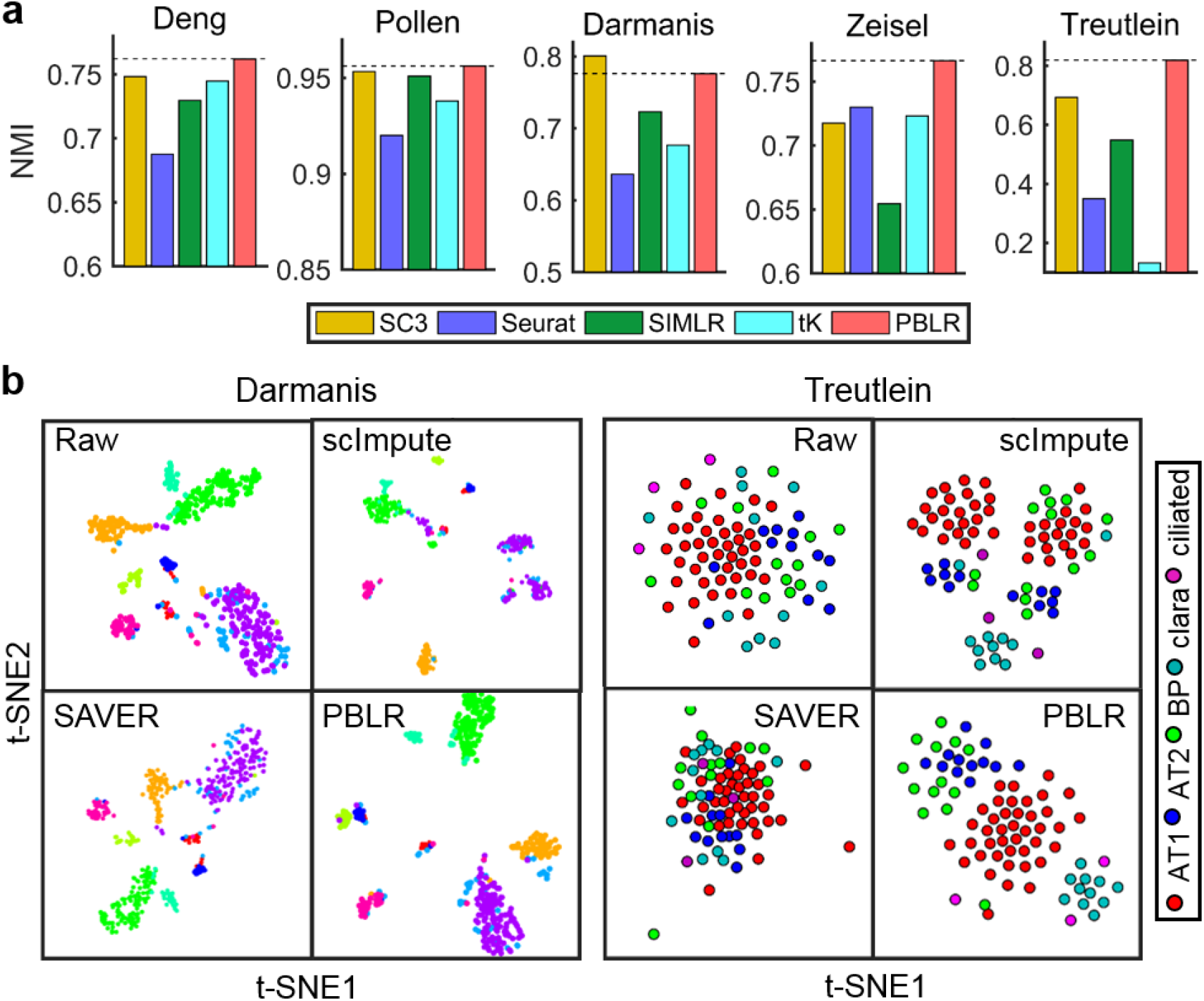
Clustering performance of PBLR and other competing methods on five real datasets. **(a)** SC3, Seurat, SIMLR, tK and PBLR were applied to the five real scRNA-seq datasets, where cell cluster labels were known or validated in the original studies. tK represents k-means on the first two t-SNE dimensions. NMI is used to quantify accuracy, **(b)** Cells are visualized by the first two t-SNE components on the raw Darmanis (left) and Treutlein (right) data, and imputed one by PBLR, scImpute and SAVER, respectively.

Moreover, visualization of Darmanis and Treutlein datasets imputed by PBLR, scImpute and SAVER or not in the first two t-SNE components demonstrates that PBLR can make various cell subpopulations more separable. AT1 and AT2 cell subpopulations are clearly distinguishable in the first two t-SNE components of PBLR imputed data. And clara cluster is separated from other ones, which is recovered by PBLR but masked by dropouts on raw data (Fig. 5b). However, other two methods either separate cells from the same cluster into several small groups (scImpute), or cannot distinguish different clusters accurately (SAVER). Therefore, PBLR can not only recover dropout events with high accuracy, but also improve precise identification of cell subpopulations compared to several state-of-the-art clustering methods.

## Discussion

We present a powerful computational method for scRNA-seq data imputation. By case studies using available scRNA-seq data from diverse investigations and synthetic data simulated with a representative tool, we demonstrate that PBLR can reduce potential dropout events and biases by considering their subpopulations and observed expression distributions, and successfully derive biologically meaningful information from data imputation. PBLR accurately recovers gene-gene relationship which is weakened by dropouts than other two competing imputation methods. Moreover, PBLR significantly improves the performance of Monocle 2 on inferring differentiation trajectory, as demonstrated in the human early embryo development and the reprogramming from mouse embryonic fibroblasts to induce neuronal cell.

As a data-driven method, PBLR uses basic principles from the low-rank matrix recovery theory by well modeling the structured information among the data. PBLR has few parameters, therefore making it more generally applicable to data from diverse labs or techniques.

PBLR consists of two key stages including identifying cell subpopulations and imputing dropouts. In the first stage, PBLR scales up well when the number of cells increases. In the second stage, singular value decomposition thresholding is the most time-consuming step. And the computational efficiency will improve if feature selection and partial singular value decomposition method being used. PBLR is an interactive method, cluster number and boundary function can be adjusted by users according to the characteristics of their datasets. Here the cluster number is selected based on clustering stability. It definitely can be used if the cluster number is known in advance in some situations.

As the high dimension of scRNA-seq data, dimension reduction is a powerful analyzing strategy. However, some meaningful low-dimensional representations are masked by dropouts. PBLR can accurately remove the influence of dropouts in low dimensions on both synthetic and real datasets. Identifying cell subpopulations is a coproduct of PBLR. Therefore, the utility of PBLR is very flexible that it can also be used to achieve a subpopulation identification task. Comparison with existing clustering methods on real datasets demonstrates that PBLR also has more accurate clustering performance.

Taking together, PBLR can be used as a general method for addressing the dropout events prevalent in scRNA-seq data with the potential to reduce noise and correct biases. It serves as a proof of principle that bias can be removed by such a classical matrix recovery methodology with more practical considerations. Moreover, PBLR can be extended to impute data for other single-cell omics data by adapting its practical boundary observations. It provides a novel approach to omics data imputation, an area that is becoming increasingly important for improving big biological data in the single-cell biology era.

## Methods

### Datasets and preprocessing

The simulated datasets were generated by Splatter^20^, an R package used for simulating scRNA-seq data. The parameters used to generate synthetic datasets 1-3 are shown in Supplementary Table 1.

We adopted two real datasets in this study for exploring expression dynamics. HEE dataset^21^ is a single cell gene expression data consisting of 88 cells from seven stages (from oocytes to blastocyst) during human early embryo (HEE) development. Finally, we obtained a data matrix with 16658 genes across 88 cells after filtering out genes expressed in less than 5 cells. MEF dataset^23^ was used to dissect the reprogramming from mouse embryonic fibroblasts (MEFs) to induce neuronal (iN) cells. To reconstruct the reprogramming path from MEFs to iN cells, similar to the original study^23^, we used 221 cells collected at multiple time points (0, 2, 5, 22 days) after removing cells that appeared stalled in reprogramming due to Ascl1 silencing or cells converging on the alternative myogenic fate.

We adopted five real datasets in this study for cell subpopulation identification (Supplementary Table 3). Deng dataset^27^ consists of 22431 genes across 268 cells, which were taken from the mouse embryo development process from zygote to blastocyst. Pollen dataset^28^ contains 301 single cells across diverse tissues, including neural cells and blood cells. It was used to test the utility of low-coverage scRNA-seq to identify cell subpopulations. Darmanis dataset^29^ was used to capture the cellular complexity of the adult and fetal human brain, including 20214 genes across 90 cells. These cells were divided into six groups, including astrocytes, endothelial, microglia, neurons, fetal quiescent and fetal replicating. Zeisel dataset^30^ contains 3005 single cells came from mouse cortex and hippocampus. The cells were collected by unique molecule identifier (UMI) and divided into nine clusters. Treutlein dataset^31^ was taken from distal mouse lung epithelial cells at different developmental stages. We used 80 single cells at E18.5 stage, which were clustered into five groups including BP, AT1, AT2, Clara and Ciliated.

For each dataset, genes expressed in less than 3 cells (unless noted specifically) and cells with expressed genes less than 200 were removed. Then the data was normalized by a global method, i.e., expression of each gene was divided by the total expression for each cell, multiplied a scale factor (10,000 by default) and log-transformed with pseudo-count 1.

### Gene selection

To account for technical noise in scRNA-seq data and select the informative genes, a set of highly variable genes were identified by calculating the average expression and Fano factor for each gene, binning the average expression of all genes into 20 evenly sized groups and then normalizing the Fano factor^32^ within each bin. Genes with a larger normalized Fano factor value (0.05 by default) and its average expression being in predefined range (0.01 to 3.5 by default) were selected. Moreover, genes with larger Gini index values^16,17^ can also be helpful to identify rare cell subpopulations (as used in Treutlein dataset).

### Affinity matrices calculation

The distance between each cell pair was computed by Pearson, Spearman and Cosine metrics, respectively. These distance matrices (denoted by *D_k_*) were transformed to affinity matrices as follows: 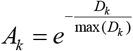.

### Subpopulation and submatrix determination

We first adopt symmetric non-negative matrix factorization (SymNMF)^33^ and incomplete non-negative matrix factorization (INMF) to the affinity matrices and raw scRNA-seq data to determine the consensus map, respectively. (1) SymNMF decomposes a non-negative affinity matrix into two symmetric non-negative low-rank matrices as follows,

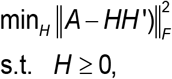

where *A* is the affinity matrix and *H* is the non-negative low-rank matrix, which can be used to indicate clustering assignment. As SymNMF is a non-convex problem that may lead to the assignment being not unique, we repeat it 20 times with random initial values. (2) Let *M_s_* represent the raw expression matrix with selected genes as its rows and cells as its columns. Let S represent the indicator matrix with element *S(i,j)*=1 if *M_s_(i,j)* is a non-zero value, otherwise *S(i,j)*=0. The following INMF model is used to learn a low-rank coefficient matrix *H_s_* to assign each cell to one cluster,

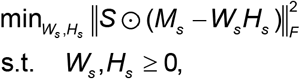

where ☉ is dot product. Similar to SymNMF, we also repeat INMF 20 times with random initial values. SymNMF and INMF are solved by alternative nonnegative least square and multiplier update algorithm, respectively.

Here, we adopt a consensus clustering method^34^ to identify cell subpopulations of cells. Each column’s maximum value of *H* or *H_s_* obtained from SymNMF or INMF under each run is used to determine the cluster membership^35^. The membership can be represented by a connectivity matrix *C*, with element *C(i,j)* = 1 if cell *i* and cell *j* are assigned into the same cluster, otherwise *C(i,j) = 0*. Then the connectivity matrices are summed across all runs and normalized by the number of runs. Thus, we obtain a consensus matrix 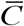 and the entries vary from 0 to 1. The entry represents the probability of cells being grouped together. Next, hierarchical clustering (HC) with average linkage is applied on 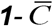, where ***1*** is matrix with all entries equaling 1. The clustering stability can be estimated by the cophenetic correlation coefficient *ρ*, which is computed as the Pearson correlation of 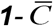 and the distance between cells inferred by average linkage. Let *ρ*_1_ represent coefficient obtained from the average consensus matrix of Pearson, Spearman and Cosin distance, and *ρ*_2_ stands for that from the consensus matrix computed from INMF. If |*ρ*_1_ − *ρ*_2_| > *cutoff*, the final clustering result is computed by the average linkage HC on 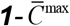, where 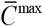 means the consensus matrix of the lager coefficient. If|*ρ*_1_ − *ρ*_2_| < *cutoff*, the final clustering result is computed on 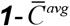, where 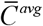 is the average of all consensus matrices. Finally, we get *g* cell subpopulations of cells, and *g*+1 corresponding submatrices *(M^(k)^*, *k*=1, …, *g*+1) of the raw scRNA-seq data *M* by extracting the sub-matrix *M^(k)^* (*k*=1, …, *g*) of each cell population of selected genes, and sub-matrix *M^(g+1)^* of the remaining genes across all cells. An optimal low rank k can be selected from a given range with the stability of clustering associated with each rank^34^. We select values of *k* where the magnitude of 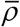 begins to fall, where 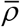 is computed by the Pearson correlation of 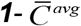 and the distance between cells inferred by average linkage on 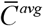.

### Boundary estimation

Let *M^(k)^* represent the *k*-th submatrix of gene expression matrix. We first compute the average expression *g_i_* of gene *i* in the observed space and the ratio of zeros *r_i_*. We only use the genes with *r_i_* being not equal to 0 and 1 because these genes either have no dropout (i.e., *r_i_*= 0) or are not expressed in all cells (i.e., *r_i_* = 1). After removing these genes, we estimate the upper boundary of gene *i* in the following ways. One way is to fit the ratio of zeros *r* versus average expression level *g* with *r* = *e*^−*λg*^2^^, then the boundary of each gene is defined as the upper one-sided 95% confidence bound. However, we find that this exponential function does not fit well for some larger *r* and overestimate the boundary (Supplementary Fig. 1). Therefore, we attempt to determine the boundary of gene *i* by introducing a piecewise function *U_i_*. First, to estimate the boundary of gene *i*, we define its neighbor gene set *S* = {*j* | |*r_j_*-*r_i_*|<*c*) using a radius *c* (default 0.05). Then, we compute the boundary of gene *i* by

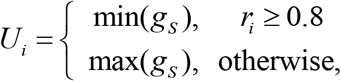

where *g_s_* = {*g_j_* | *j* ∈ *S*} is the expression of the neighbor gene set. Moreover, we define a more sophisticated piecewise function,

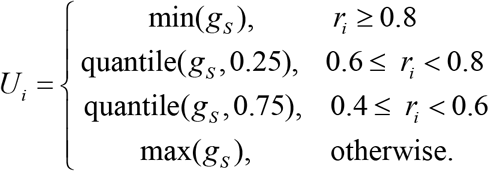

The sophisticated piecewise function is used as default (see Supplementary Note). However, we also recommend choosing a proper boundary function by visually evaluating the scatter plot of ratio of zeros versus average expression level on a sampled reference data (Supplementary Fig. 1). We generated a reference data by dropping varying fractions (relevant to the dropout rate) of the gene measurements in the raw gene expression matrix. We simulated dropouts by setting true values to zero by sampling from a Bernoulli distribution using a dropout probability max(*p*_0_,0.3), where *p*_0_ is the ratio of zeros in the raw expression matrix.

### Bounded low-rank imputation algorithm

We adopt an ADMM algorithm^18,19^ to solve the bounded low-rank matrix recovery model. Specifically, it can be reformulated as follows,

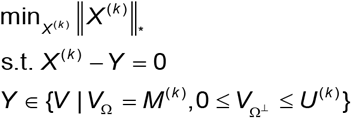

The augmented Lagrangian function of the above function is

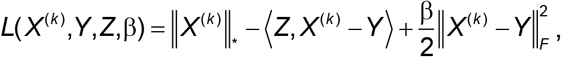

where *Z* is the Lagrange multiplier, β is the penalty parameter. We update the variables by alternatively updating *X^(k)^*, *Y*, *Z* as follows,

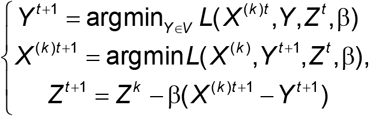

where *t* is the iteration index. In more detail, we can update variable *Y* by argmin*_Y_* 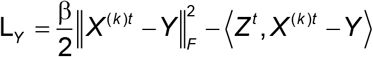.

Note that the partial derivative on *Y* of *L_Y_* is equal to *Z^t^ − β*(*X*^(*k*)*t*^ − *Y^t^*), therefore, it can be reformulated as 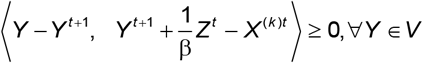. The solution is 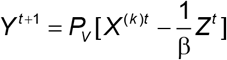, where *P_V_* is the projection operator onto *V* space. The solution can be written as follows,

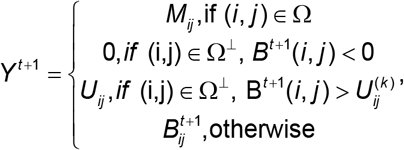

where 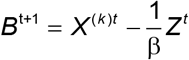. Then let 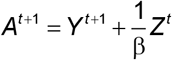, and 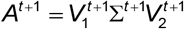, where 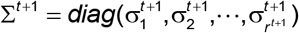 and 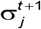 is the eigenvalues of *A*^*t*+1^. According to a traditional solution in previous studies^36,37^, the update rule for *X* is 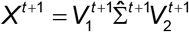, where 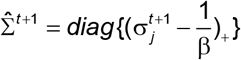. Therefore, we only need to compute the eigenvalues larger than 1/*β* and we use PROPACK package to compute the partial SVD. Previous studies^38,39^ have proved that the step for updating the Lagrange multiplier can be generalized into 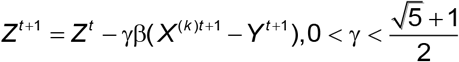. In the proposed algorithm, we use the same parameter γ = 1.6 and 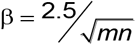 as in a previous study^18^. This procedure is summarized in **Algorithm 1**.

#### Algorithm 1 BLR

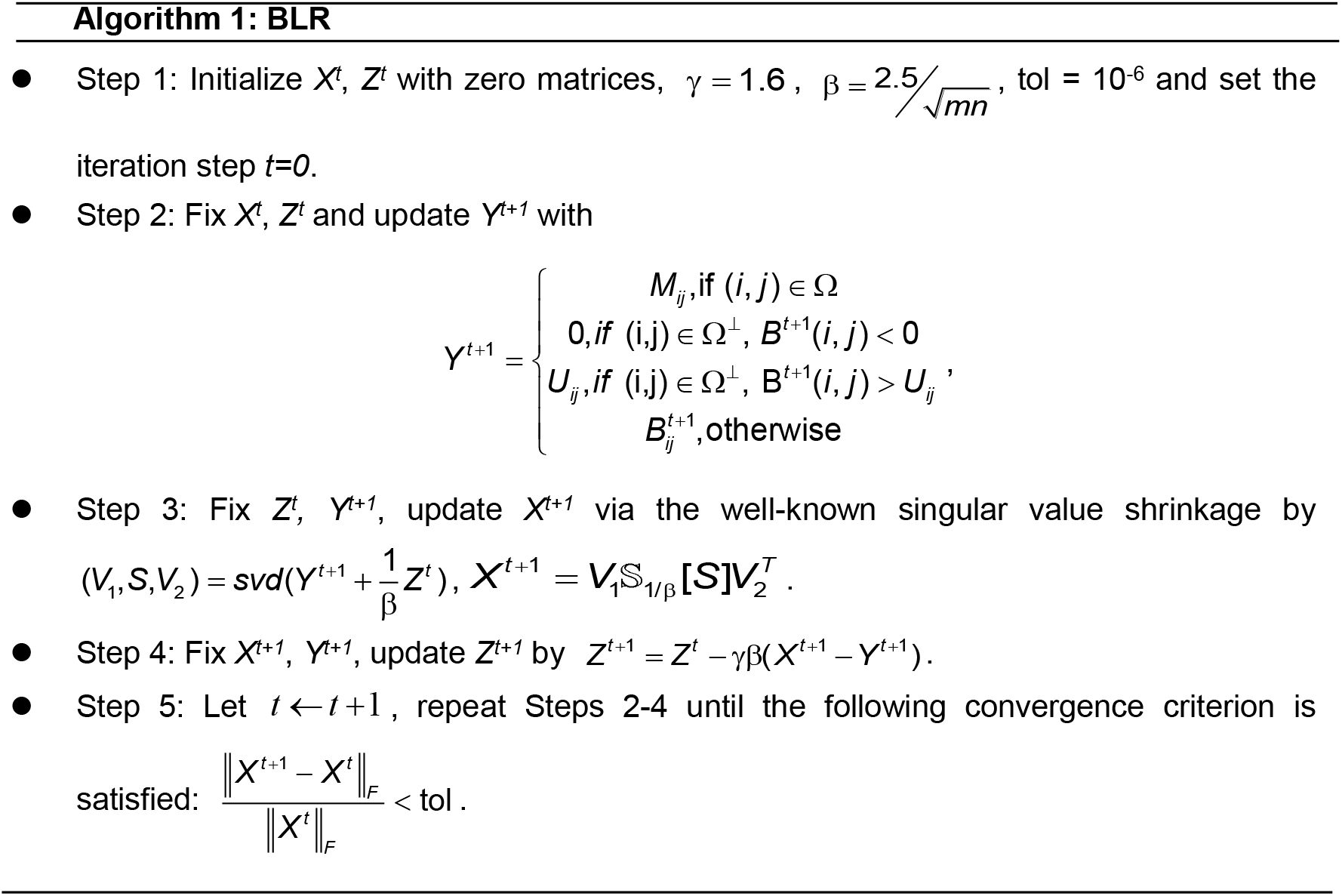

### PBLR algorithm

The whole procedure for solving scRNA-seq imputation is summarized in **Algorithm 2**.

#### Algorithm 2 PBLR

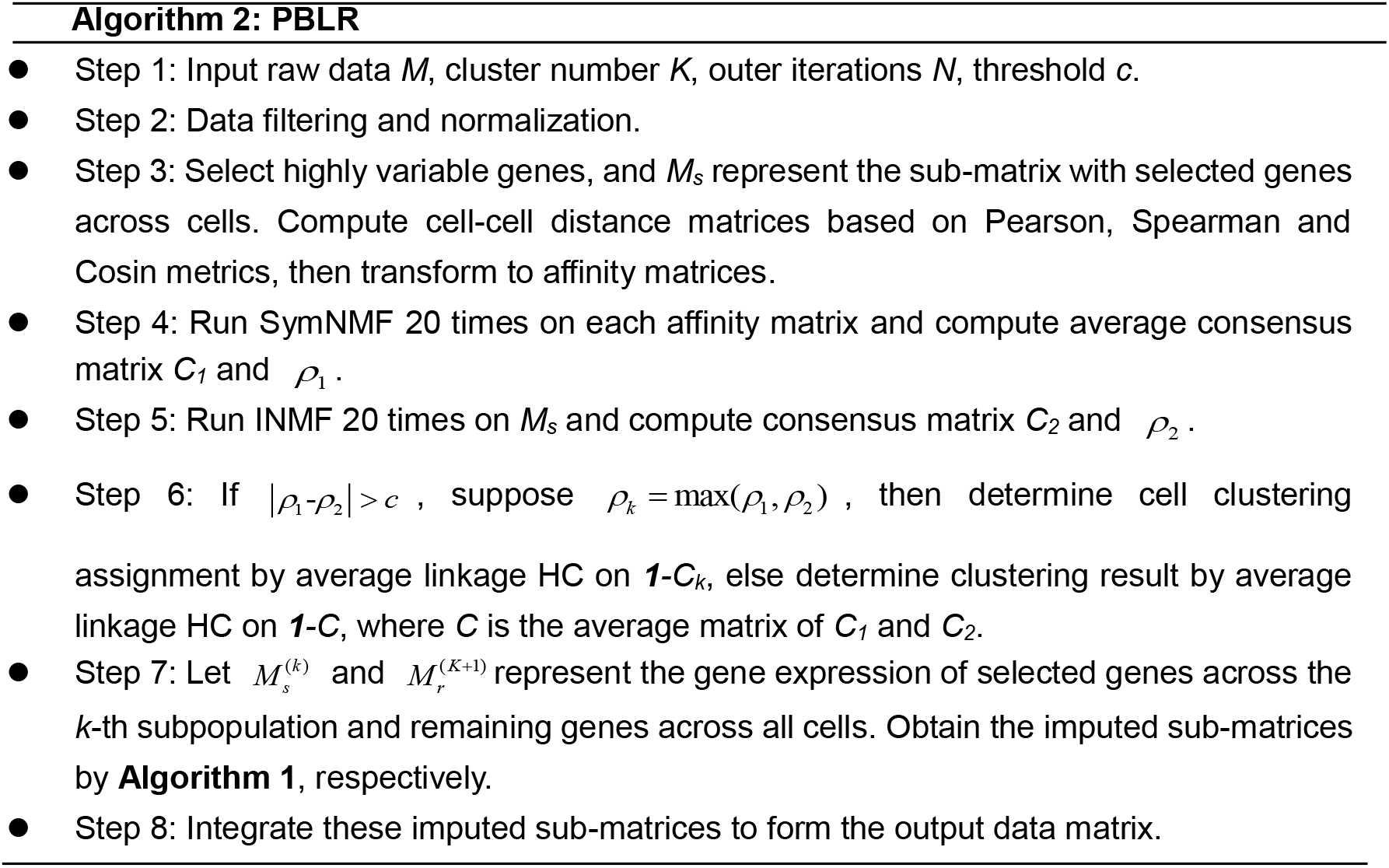

### Imputation accuracy evaluation on synthetic datasets

To quantify the difference between imputed data and full data, we calculated two measures: sum of squared error (SSE) and Pearson correlation coefficients (PCC). SSE is defined as 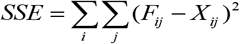, where *F_ij_* represents the real expression of gene *i* in cell *j*, while *X_ij_* represents the corresponding imputed value. PCC is computed between each column pair (*F_.j_* and *X_.j_*) of *F* and *X*.

### Normalized mutual information (NMI)

We use *U* = {*U*_1_, …, *U_m_*} to denote the true partition of *m* classes and *V* = {*V*_1_, …, *V_n_*} to denote the partition given by PBLR. Then 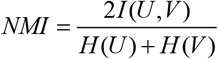, where *I(U,V)* is mutual information, *H(U)* is the entropy of partition *U*.

### Pseudotime order score (POS)

To measure the accuracy of the reconstructed pseudotime, we define a pseudotime order score (POS): POS = *C/(Nc* + *C)*, where *C* and *N_c_* represent the number of concordant and disconcordant pairs of cells between the inferred pseudotime and golden standard (e.g. true data collection time), respectively.

### Code availability

PBLR is written as a Matlab package which is available at http://page.amss.ac.cn/shihua.zhang/software.html.

### Data availability

Deng, Darmanis, Treutlein and Zeisel datasets can be obtained from Gene Expression Omnibus (GEO) with GSE45719, GSE6785, GSE52583 and GSE60361 respectively. Pollen dataset is available at Sequence Read Archive (SRA) with SRP041736. HEE and MEF datasets can be obtained from GEO with GSE36552 and GSE67310 respectively.

